# Ultrastructural analysis reveals mitochondrial placement independent of synapse placement in fine caliber C. elegans neurons

**DOI:** 10.1101/2023.05.30.542959

**Authors:** Danielle V. Riboul, Sarah Crill, Carlos D. Oliva, Maria Gabriela Restifo, Reggie Joseph, Kerdes Joseph, Ken C.Q. Nguyen, David H. Hall, Yaouen Fily, Gregory T. Macleod

## Abstract

Neurons rely on mitochondria for an efficient supply of ATP and other metabolites. However, while neurons are highly elongated, mitochondria are discrete and limited in number. Due to the slow rates of diffusion over long distances it follows that neurons would benefit from an ability to control the distribution of mitochondria to sites of high metabolic activity, such as synapses. It is assumed that neurons’ possess this capacity, but ultrastructural data over substantial portions of a neuron’s extent that would allow for tests of such hypotheses are scarce. Here, we mined the *Caenorhabditis elegans’* electron micrographs of John White and Sydney Brenner and found systematic differences in average mitochondrial length (ranging from 1.3 to 2.4 μm), volume density (3.7% to 6.5%) and diameter (0.18 to 0.24 μm) between neurons of different neurotransmitter type and function, but found limited differences in mitochondrial morphometrics between axons and dendrites of the same neurons. Analyses of distance intervals found mitochondria to be distributed randomly with respect to presynaptic specializations, and an indication that mitochondria were displaced from postsynaptic specializations. Presynaptic specializations were primarily localized to varicosities, but mitochondria were no more likely to be found in synaptic varicosities than non-synaptic varicosities. Consistently, mitochondrial volume density was no greater in varicosities with synapses. Therefore, beyond the capacity to disperse mitochondria throughout their length, at least in *C. elegans*, fine caliber neurons manifest limited *sub-*cellular control of mitochondrial size and distribution.

**SIGNIFICANCE:** Brain function is unequivocally reliant on mitochondrial function for its energy needs, and the mechanisms that cells use to control these organelles is an active field of enquiry. WormImage, a decades old electron microscopy database in the public domain, contains information about the ultrastructural disposition of mitochondria within the nervous system of *C elegans* over previously unexamined extents. In a largely remote format, a team of students mined this database over the course of the pandemic. They found differences in mitochondrial size and density between neurons, but limited differences between different compartments of the same neurons. Also, while neurons are clearly able to disperse mitochondria throughout their extent, they found little evidence that they “install” mitochondria at synaptic varicosities.

## INTRODUCTION

Neurons are highly elongated, allowing them to communicate over long distances, but their morphology presents challenges for small outposts with high metabolic demands, such as synaptic varicosities. Mitochondria are essential components of neuronal metabolism, and, among other functions, they generate ATP, buffer Ca^2+^ and contribute to neurotransmitter synthesis (Devine and Kittler, 2018). However, due to the limitations to diffusion of both mitochondria and metabolites, neurons need mechanisms to move mitochondria to, and/or concentrate them, where they are most needed (Cheng et al., 2022; Lopez-Domenech and Kittler, 2023). Motor-based transport distributes mitochondria along microtubules and microfilaments to a neuron’s furthest extent (Kruppa and Buss, 2021), while other mechanisms can arrest their movement or anchor them in place. For example, posttranslational modification of microtubules modifies motor-based transport (Hammond et al., 2010), local elevations in Ca^2+^ arrest mitochondrial movement through the agency of the adaptor protein *miro* (Wang and Schwarz, 2009), *syntaphilin* and FHL2 anchor mitochondria along axons (Kang et al., 2008; Basu et al., 2021), VAPB anchors mitochondria along dendrites (Bapat et al., 2024), and membrane interactions arrest mitochondrial adjacent to other organelles (Lebiedzinska et al., 2009). This list is far from exhaustive, but it demonstrates numerous mechanisms available to modify motor-based dissemination of mitochondria, presumably to position them where they are most needed.

Our understanding that neurons distribute mitochondria to synapses, and exert subcellular control of mitochondrial morphology and volume density, is based on fluorescence light microscopy data and electron microscopy data. However, limitations of both techniques leave room for reinterpretation of the data. The spatial resolution of light microscopy is inadequate for the purposes of estimating mitochondrial number, morphology or volume density, especially when they are densely intertwined. Electron microscopy, while providing exceptional resolution, has a greatly limited field of view, which generally excludes the opportunity to sample neurons in their entirety, or to sample ultrastructural features in sufficient numbers for statistical analyses. There are notable exceptions. Super-resolution techniques have increased light microscopy resolution of mitochondria and synapses by an order of magnitude (Huang et al., 2008; Sigal et al., 2015), but the fields of view have shrunk commensurately. At the same time, automation of aspects of the electron microscopy work flow have allowed for the reconstruction of greater volumes of nervous tissues (yet still < 1% of 1 cubic millimeter) in both mammals and invertebrates (Turner et al., 2022; Winding et al., 2023), although these data repositories have yet to reveal the disposition of individual mitochondria relative to each other, or relative to synapses. Analysis likely trails the acquisition of the data because of the difficulty in automating and validating protocols to quantify distances and volumes between mitochondria and synapses within the organic shapes of neuronal processes.

In this study, analysis has trailed data acquisition by 40 years. We reconstructed motor neurons, interneurons and sensory neurons from transmission electron micrographs spanning a 442 μm length of *C. elegans* (White et al., 1986). The unambiguous identification of each neuron, high quality of mitochondrial profiles, recent revision of synapse annotation (Cook et al., 2019) and the relatively simple morphology of these neurons gave an unprecedented opportunity to interrogate the disposition of individual mitochondria relative to each other and to synapses. We observed differences in mitochondrial morphometrics between neuron types, but we found limited evidence for either subcellular control of mitochondrial volume density or mechanisms that control mitochondrial approach to synapses. While neurons in *C. elegans* generally do not rely on classical all-or-none Na^+^-based action potentials (Lockery and Goodman, 2009), these data never-the-less raise questions about whether we should anticipate similarly limited mitochondrial control in fine caliber neurons in vertebrates *in vivo*.

## RESULTS

### Neurons can be reconstructed in their entirety from WormImage electron micrographs

Transmission electron micrographs collected from the *C elegans* N2U section series allowed us to reconstruct neurons over a substantial extent of their length. We examined dorsal micrographs labeled #1 to 1763 and ventral micrographs #1 to 1844 (Fig. 1A). Sequentially labeled micrographs posterior to the esophageal ganglion represent every 3^rd^ section of a larger series of sections cut at 80 nm intervals (Cook et al., 2019). Using ImageJ software we measured the cross-sectional areas of neuronal profiles, and, separately, the mitochondria within (Fig. 1A). Seventeen (17) neurons were selected to represent a manageable sample of the neurotransmitter and functional diversity, yet still providing repetition across 5 neuron types; GABAergic motor neurons (5), cholinergic motor neurons primarily innervating body wall muscles (3), cholinergic motor neurons primarily innervating vulval muscles (3), cholinergic interneurons (4) and glutamatergic sensory neurons (2). Each neuron is represented in silhouette where its diameter was calculated from the area of the neuronal profile rendered as a perfect circle (Fig. 1B). The GABAergic motor neurons have both dorsal and ventral processes connected by commissures (commissures not shown). The series covered between 28 and 100% of the longitudinal extent of 17 neurons, (73 ± 28%; mean ± SD), capturing the somata of all 11 motor neurons (Fig. 1B).

**Figure 1.**
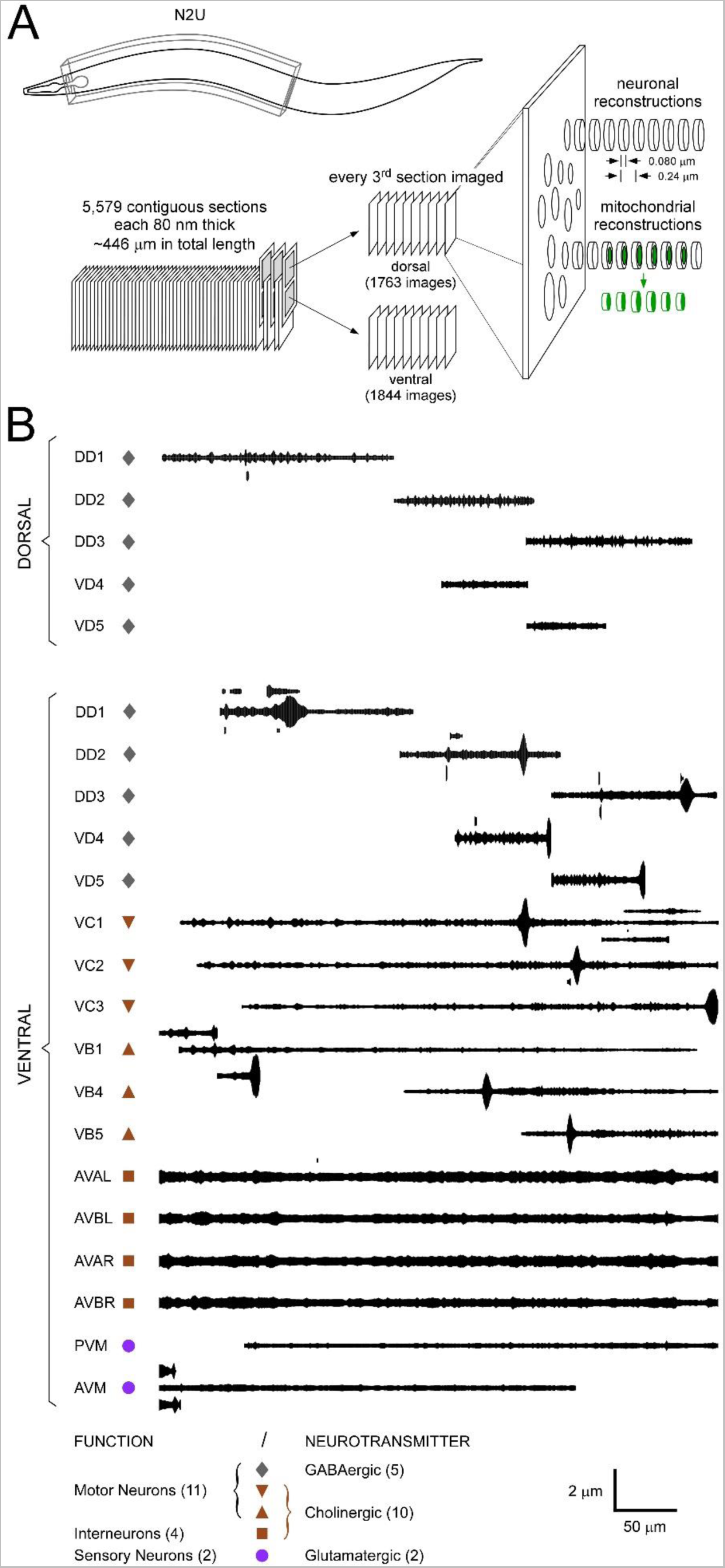
3D reconstruction of 17 neurons from an electron micrograph series through a worm. (A). Schematic of the N2U series in *C elegans* and the workflow from contiguous micrographs to reconstructions using neuronal and mitochondrial cross-sectional areas measured in ImageJ. The series spans the anterior portion of a hermaphroditic worm and is composed of 5579 sections, each 80 nm thick. Photomicrographs were taken of every 3^rd^ section covering a portion between the posterior extent of the nerve ring and the vulva. We analyzed 1844 contiguous micrographs on the ventral side, posterior to the nerve ring and neuropile. 1763 contiguous micrographs were analyzed on the dorsal side. Each area measurement represents the average for a section, separated from the next by 160 nm (240 nm between centers). (B). Silhouette reconstructions of 17 neurons. Profiles smoothed with a 5-point moving average. A legend for the symbols beside individual neuron names is shown at the bottom. Individual GABAergic motor neurons (DD1, DD2, DD3, VD4, VD5) have a dorsal and a ventral extent. Enlarged swellings indicate cell bodies captured in this micrograph series for some of the neurons (DD1, DD2, DD3, VD4, VD5, VC1, VC2, VC3, VB1, VB4, VB5). Branches exceeding 1 μm in length are represented on one (VD4, VC2), or both (DD3, VC1, VB1), sides of the primary axis of some neurons. Anterior to posterior: left to right. Name acronyms: Dorsal D-type Motor Neuron 3 (DD1, DD2, DD3); Ventral D-type Motor Neurons 4 and 5 (VD4 and VD5); Ventral B-type Motor Neurons 1, 4 and 5 (VB1, VB4 and VB5); Ventral C-type Motor Neuron 1, 2 and 3 (VC1, VC2 and VC3); Anterior Ventral Processes A Left and Right (AVAL and AVAR); Anterior Ventral Processes B Left and Right (AVBL and AVBR).

### The lateral extent of mitochondria can be resolved at 5 nm resolution

The N2U series was originally photographed by (White et al., 1986) and our measurements were made on scanned images of these micrographs, digitized in a range between 140 – 840 pixels / micron (Fig. S1). Mitochondrial profiles could be resolved with strong contrast, often allowing discrimination of the outer membrane separate from the inner membrane and showing cristae structure (Fig. 2A-E, panels on left). Neurons in each micrograph are annotated with a handwritten identifying number (Fig.2A-E, panels on left), and the extent of each mitochondrion is represented within each neuron, centered on the long axis (Fig.2A-E, panels on right).

**Figure 2.**
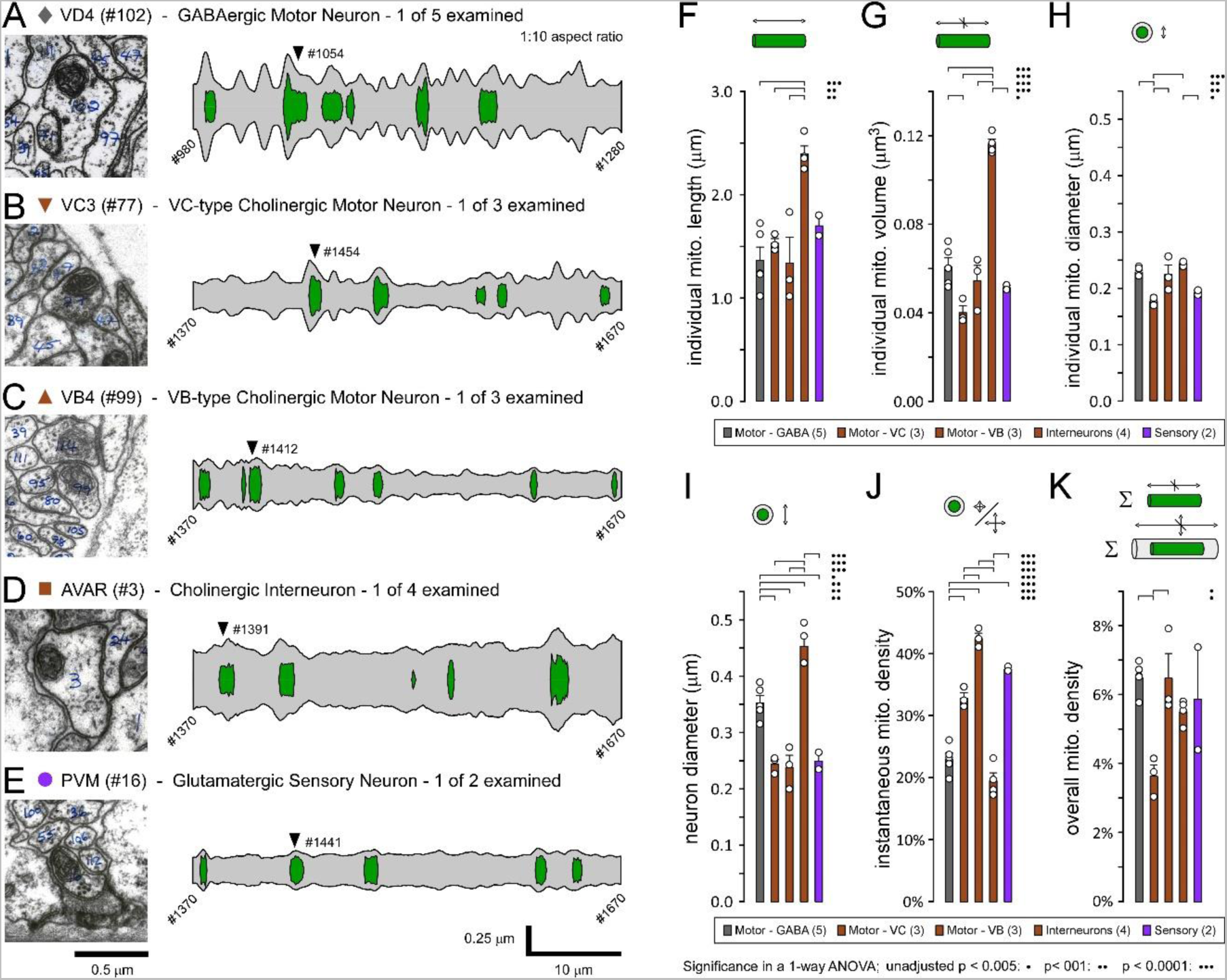
Mitochondrial morphometrics differ according to neuronal function. A-E. 2D reconstructions of example neurons from different 5 classes, along with their mitochondria. Left panels: 1 μm x 1 μm regions of interest cropped from single micrographs in the N2U series. Right panels: neurons and mitochondria reconstructed from cross-sectional area measurements made in each of 301 micrographs. Profiles smoothed with a 5-point moving average. Neuron identity and neurotransmitter type indicated above each micrograph in A-E. Arrowheads show the number and location of each example micrograph. F-K. Plots comparing mitochondrial and neuronal morphometrics between 17 neurons divided into their functional and neurotransmitter type. F. For each neuron, individual mitochondrial length was calculated by multiplying the number of sections by 0.24 μm, and then subtracting 0.08 μm from the resulting total (see Methods). G. Individual mitochondrial volume was calculated by multiplying the average mitochondrial area by the length of the mitochondrion. H. Individual mitochondria diameter was calculated by dividing the average mitochondria area by pi (π), then taking the square root and multiplying by 2. I. Neuronal diameter was calculated by dividing the average neuronal profile area by π, then taking the square root and multiplying by 2. J. Instantaneous mitochondrial density was calculated by dividing summed mitochondrial areas by the neuronal profile area (the latter including both cytosolic and mitochondrial areas) across all micrographs where the neuron contained a mitochondrion. K. Overall mitochondrial density was calculated by dividing summed mitochondrial areas by summed neuronal profile areas. Areas were calibrated according to the number of pixels that spanned the known distance between thick filaments in the cross-sectioned muscle. See methods and materials for further details. One-way ANOVAs, with Holm Sidak post hoc tests, show significance at four levels: <0.005 (1 dot), <0.001 (2 dots), <0.0001 (3 dots). α = 0.005 represents an α = 0.05 Bonferroni correction across 10 tests.

### Mitochondrial diameter and density differ systematically between the five neuron types

Parameters of mitochondrial size and shape were analyzed in both neuronal cell bodies (somata) and processes, but only data for processes are shown in figure 2. Average mitochondrial length and volume were calculated for each of the 5 neuron types and mitochondria were found to be twice as large in cholinergic interneurons (length and volume; Fig. 2F & 2G). Average mitochondrial diameters have a tight variance and appear to be under exquisite control, with mitochondria in VC-type cholinergic motor neurons having the “thinnest” mitochondria (Fig. 2H). Average neuronal diameter is also tightly controlled, with command interneurons and GABAergic motor neurons having the largest calibers (Fig. 2I). A plot of mitochondrial diameter against mitochondrial length showed no association indicating that mechanisms determining length and diameter can work independently (Fig. S2A; P=0.541). Mitochondrial diameter is necessarily constrained by neuron diameter, and an association between the two was observed in a smaller organism – a miniature parasitic wasp (Fischer et al., 2018). Here, however, we found no evidence for an association between mitochondrial diameter and neuronal diameter, thus ruling out mitochondrial diameter as a simple function of neuron diameter in *C elegans* (Fig. S2B; P=0.142). Instantaneous mitochondrial density, i.e. that proportion of a neuron’s cross-sectional area occupied by a mitochondrion, was distinct in all 5 neuron types, with VB-type motor neurons having double the instantaneous density of interneurons, despite releasing a common neurotransmitter; acetylcholine (Fig. 2J). Finally, overall mitochondrial density calculations showed VC-type cholinergic motor neurons to have a density of only 3.7%, while the other 4 neuron types grouped in a range with almost twice the density (between 5.5% and 6.5%; Fig. 2K).

### Limited subcellular control of mitochondrial length, size and density

In an initial test of the neurons’ capacity for subcellular control of mitochondrial morphology and volume density we compared parameters measured in Figure 2F, 2H, 2I & 2K between different neuronal compartments. Each neuron might be divided into different compartments according to which portion is in the ventral versus dorsal nerve cord, which portions are cellular processes versus the soma, and which portions are primarily dendritic versus axonal. The latter distinction was made by annotating each neuron profile with the micrograph/section location of either presynaptic (blue section) or postsynaptic specializations (pink section) while cross checking with Cook and others (2019) (Fig. 3A-C).

**Figure 3.**
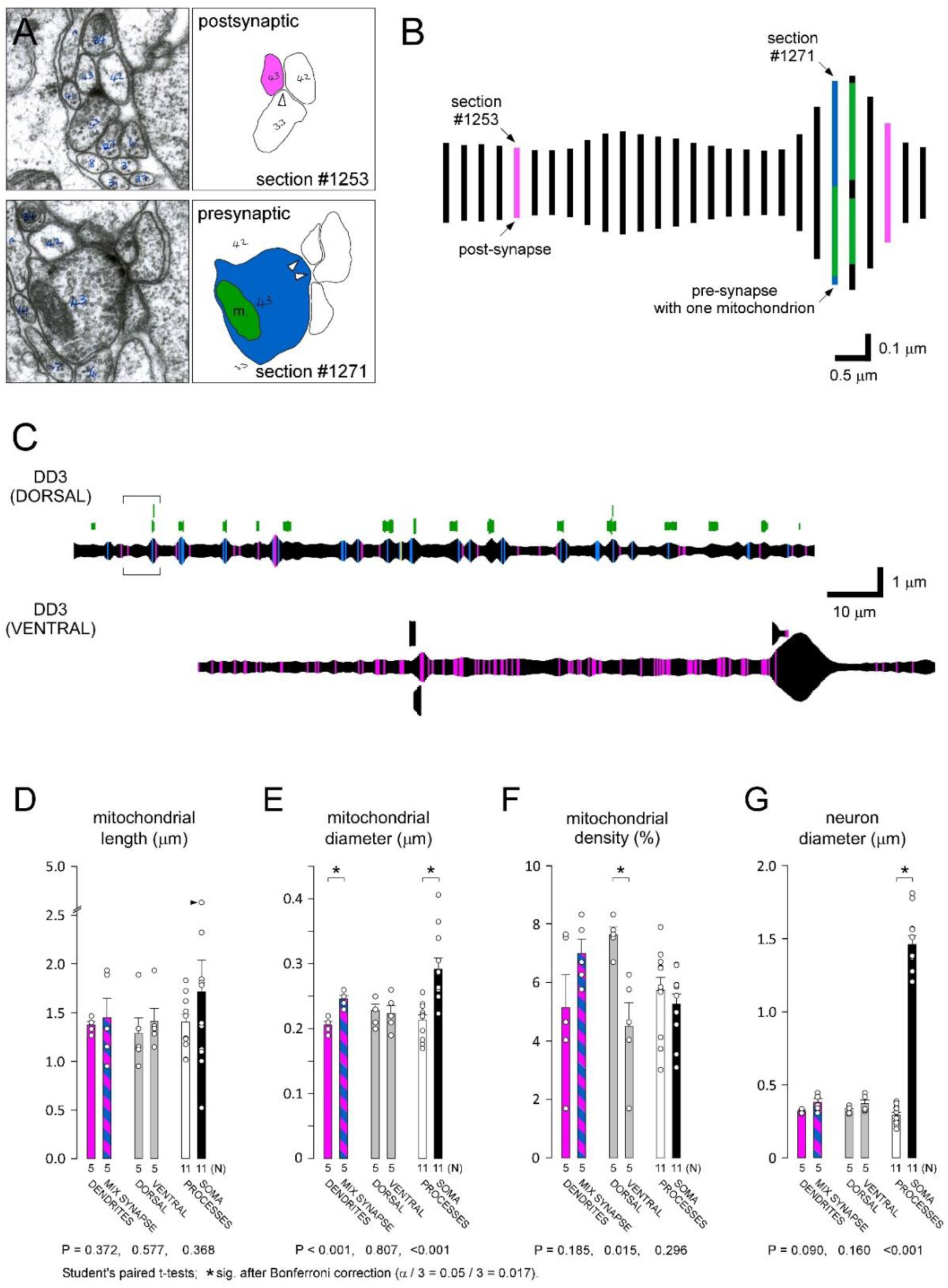
Mitochondrial morphometrics show few differences across subcellular compartments in motor neurons. A. Upper image: micrograph of section 1253 where GABAergic motor neuron DD3 is post synaptic to ventral cord motor neuron DB3. Lower image: micrograph of section 1271 where GABAergic motor neuron DD3 is presynaptic to dBWMR11 (body wall muscle); mitochondria indicated in green. B. Schematic of dorsal portion of GABAergic motor neuron DD3 from 1249-1276. Locations of pre and post synapse micrographs illustrated in A. Mitochondria indicated in green. C. Reconstruction of entire GABAergic DD3 motor neuron (excluding the commissure). Post-synapses indicated in pink, pre-synapses indicated in blue, mitochondria indicated in green (dorsal portion of DD3 only). The large swelling indicates the cell body and short processes indicate branches. D. E. F. G. Plots comparing mitochondria length, diameter, density and neuron diameter between different subcellular portions of neurons; portions containing post-synapses only (dendrites) versus portions containing both pre- and post-synaptic elements (mixed), dorsal versus ventral; and neuronal processes versus the soma. Calculations for length, diameter, density are the same as in figure 2. Paired student t-test with Bonferroni alpha correction for 3 tests (α = 0.05 / 3 = 0.017); for 12 tests (α = 0.05 / 12 = 0.004).

Each of the five GABAergic motor neurons can be divided into a “dendritic” potion containing only post-synaptic elements, and a “mixed” portion containing both pre- and post-synaptic elements. In a comparison of the mitochondria in these different portions, neither mitochondrial length nor volume density registered as significantly different (P=0.372 and P=0.185, respectively; Fig. 3D-F). Mitochondrial diameter, however, was ∼20% less in the dendritic portions (P<0.001; Fig. 3E). Turning to the dorsal versus ventral portions of these motor neurons, no differences were detected in mitochondrial length nor diameter (P=0.577 and P=0.807, respectively; Fig. 3D-E), but mitochondrial density was greater dorsally - furthest from the ventral cell bodies (P=0.015; Fig. 3F). This observation is consistent with a study in pyramidal cells of the mouse primary visual cortex where mitochondrial volume density increases in dendrites with distance from the soma (Turner et al., 2022). However, while the significance of the difference observed here persisted after a Bonferroni correction of α/3 (0.05/3 = 0.017), the significance would be extinguished after a more stringent correction of α across the 12 t-tests in figure 3 (0.05/12 = 0.004). The significance of the difference in mitochondrial diameter remains.

Finally, in a comparison between mitochondria in processes versus the somata of 11 motor neurons, we observed that mitochondrial diameter, but not length, registered as significantly different (P<0.001 and P=0.368, respectively; Fig. 3D,E). Surprisingly, mitochondrial density was no greater in processes when compared to the soma (P=0.296; Fig. 3F). This was unexpected as the nucleus excludes mitochondria, yet the volume of the nucleus is not excluded when calculating mitochondrial density in the soma. The implication is that, compared to the mitochondrial density within the cytosol of cellular processes, mitochondrial density is considerably greater in the cytoplasmic shell that surrounds the nucleus in the soma.

### Mitochondrial distribution appears random except where excluded by other mitochondria

To determine whether mechanisms position mitochondria relative to each other, rather than simply disperse mitochondria throughout the length of neurons, we analyzed the distribution of distances between the centers of mitochondria (Fig. 4A-C; interneurons shown as an example group). We would expect a frequency distribution of distances to be described by an exponential decay if mitochondria are positioned randomly. The outcome of Kolmogorov–Smirnov (KS) tests of the cumulative distributions against exponentials did not allow us to reject the likelihood that mitochondria are positioned randomly in interneurons (P=0.134), but we found clear evidence for a non-random distribution in GABAergic motor neurons (P<0.001) and sensory neurons (P=0.004) (Fig. 4D).

**Figure 4.**
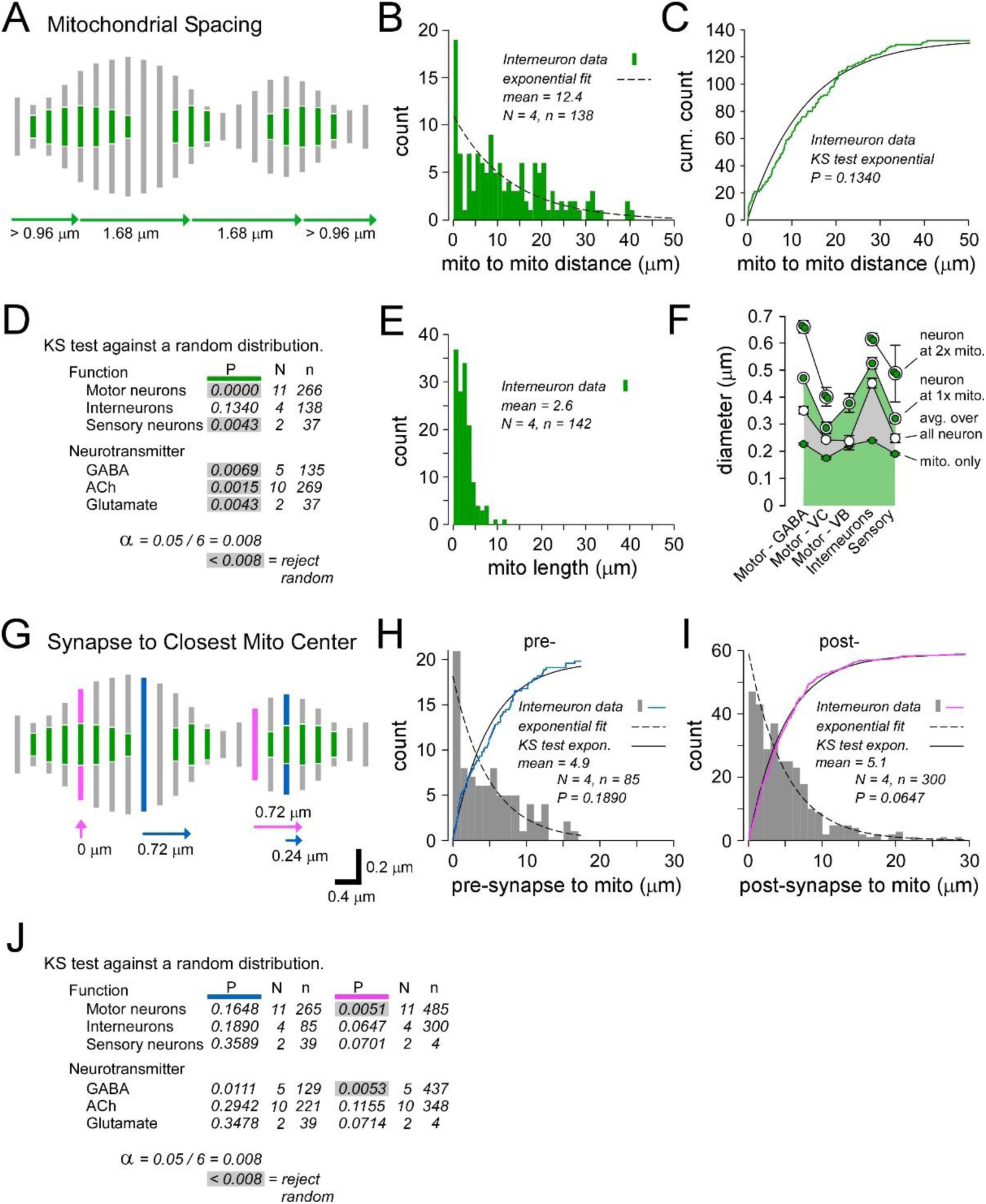
Analyses of mitochondrial positioning along fine neuronal processes. A. Schematic of a sectioned neuron with mitochondria. Distances from the center of one mitochondrion to the next mitochondrion, anterior-to-posterior, indicated with arrows. B. Frequency distributions of distances between mitochondria in cholinergic interneurons. C. A cumulative plot is shown, along with a Kolmogorov–Smirnov (KS) test against an exponential (P=0.045). The Bonferroni alpha level correction for 6 tests (panel D) is 0.008 (α = 0.05 / 6) so we are unable to reject the null that the samples are drawn from a random distribution of spacings. D. The results of KS tests against random distributions. E. A Frequency distribution of the length of mitochondria in the same cholinergic interneurons analyzed in panels B and C. F. Average diameters of mitochondria, neurons, neurons at the location of single mitochondria, and neurons at the location of overlapping mitochondria. Cell bodies were excluded. Two-way ANOVA performed on mean neuronal diameters. Neurons had a significantly greater diameter when a mitochondrion was present (P=0.006), and an even greater diameter when two were present (overlapping; P<0.0001). Neuron type factor: F(4,13)=19.8; mitochondria number present factor (0, 1 and 2): F(2,13)=38.3. G. Schematic of a sectioned neuron with AZs and mitochondria, showing the spacings that were analyzed. Pre-synapses indicated in blue, post-synapses indicated in pink and mitochondria in green. Distances from each AZ to the closest mitochondrion center are indicated with arrows. H-I. Frequency distributions of distances to mitochondria, measured from cholinergic interneuron pre- (H) and post-synapses (I). Cumulative plots are shown for each, along with KS tests against exponentials (P=0.189 and P=0.065; in both cases unable to reject the null that the samples are drawn from a random distribution of spacings). J. The results of KS tests against random distributions.

The departure from random can be readily seen in the frequency distributions for neurons grouped according to function and neurotransmitter type (Fig. S3). In each grouping, mitochondrial spacings of ∼1-5 μm are underrepresented (e.g. Fig. 4B-C). Interestingly, mitochondrial lengths form a frequency distribution (Fig. 4E) that might fill the “void” in the distribution of mitochondrial spacings. Therefore, the void in Figure 4B might be attributed to an “occlusion effect”, where the center of one mitochondrion is displaced from the center of another mitochondrion by, on average, the length of a mitochondrion, i.e. the sum of two half lengths (2.6 μm for interneurons shown in Figures 4B and E). If this is occurring, then we would expect a robust departure from random in neurons with the smallest diameters, and this is what is observed (Fig. 4D). In fine neuronal processes such as these, mitochondria appear to have little opportunity to pass one another except at varicosities. Consistently, our analysis shows that neuron diameters are not only greatest at the location of mitochondrial centers versus a random sampling of neuron diameters (Fig. 4F), but that neurons are even wider at locations where mitochondrial centers are no more than 1 μm apart, i.e. where they overlap. These analyses cannot determine causality, but the simplest explanation that fits the data is that the distribution of mitochondria is random, except where disrupted by the physical limitations to passing.

While the current study offers better than 5 nm lateral resolution in most micrographs, each micrograph represents a plan projection of 80 nm of material (section thickness) along the long axis of each neuron, with 160 nm of missing material between micrographs. The lateral resolution is therefore far superior to that offered by fluorescence microscopy (or super-resolution microscopy), but no better in the axial dimension (± 240nm). We therefore investigated the consequences of these sampling characteristics for our ability to quantify mitochondrial lengths, and the expectation that a random placement of mitochondria or synapses (or the distances between them) along the length of neurons would manifest as an exponential decay in the probability density functions of distance measurements (see Methods). Ultimately, as limitations in axial resolution were more than an order of magnitude less than the means of the exponential fits, we concluded that the limited axial resolution should not preclude the application of the Kolmogorov-Smirnov model.

### Mitochondria are randomly distributed relative to pre-synapses

To determine whether a mechanism positions mitochondria relative to pre- or post-synaptic specializations we analyzed the distribution of distances between each synapse and the center of the closest mitochondrion (Fig. 4G). KS tests of the cumulative distributions against exponentials indicated that mitochondria are generally positioned independently of pre-synapses (Fig. 4H-I), but with some support for systematic positioning of mitochondria relative to postsynaptic specializations in GABAergic motor neurons (Fig. 4J). This association should be considered in light of the analysis in figure 5F where varicosities are characterized by the presence of pre-synapses but not post-synapses, raising the possibility that a non-random association between post-synapses and mitochondria arises through the systematic exclusion of post-synapses but not mitochondria from varicosities.

**Figure 5.**
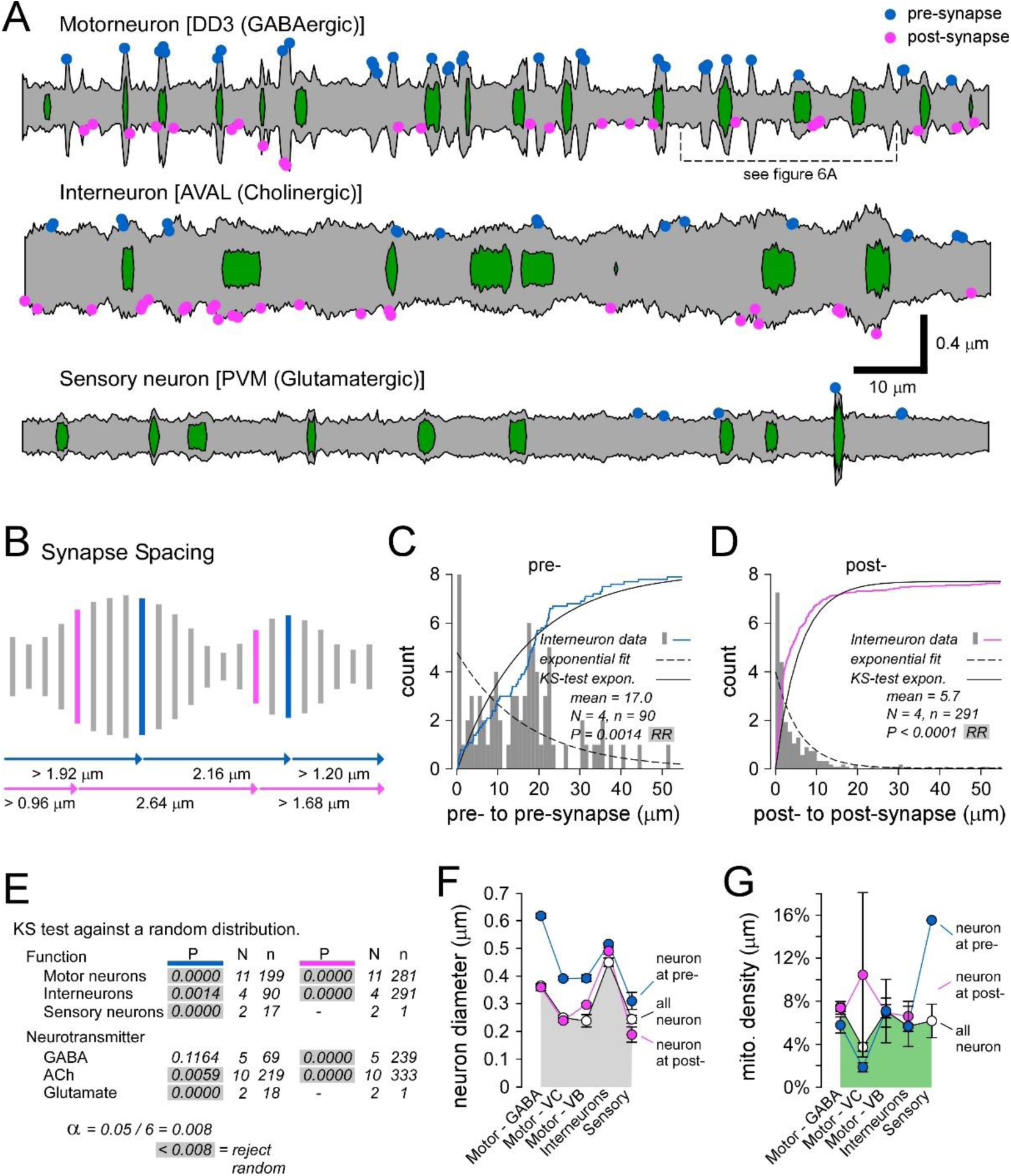
Analyses of synapse positioning along fine neuronal processes. A. Neuronal plasma membrane, mitochondria and synaptic locations, reconstructed from measurements made in each of 551 contiguous micrographs (#1213 to #1763; 132 μm; without profile smoothing). B. Schematic of a sectioned neuron showing the distances from one pre-synapse to the next pre-synapse (blue arrows), and from one post-synapse to the next post-synapse (pink arrows). C-D. Frequency distributions of distances from one synapse in a cholinergic interneuron to the next; pre- (C) and post-synapses (D). Cumulative plots are shown for each, along with Kolmogorov– Smirnov (KS) tests against exponentials (P=0.001 and P<0.0001), allowing us to reject the null that the samples are drawn from a random distribution. E. The results of Kolmogorov–Smirnov (KS) tests against random distributions. F. Average diameters of neurons, neurons at the location of a pre-synapse (blue), and at the location of post-synapses (pink). Two-way ANOVA performed on mean neuronal diameters. Neurons had a significantly greater diameter at pre-synapses (P=0.003), but not at post-synapses (P=0.822). Neuron type factor: F(4,14)=12.3; compartment identity factor (all, pre or post): F(2,14)=11.6. G. Average mitochondrial density at the location of a pre-synapse (blue), and at the location of post-synapses (pink). Two-way ANOVA performed on mean mitochondrial density. There was no significant difference in mitochondrial density at pre-synapses or post-synapses. Neuron type factor: p=0.864, F(4,12)=0.306; compartment identity factor (all, pre or post): p=0.255, F(2,12)=1.731.

### Synapses are not randomly distributed

Next, we turned our attention to the spacing between synapses (Fig. 5A-B). Our analysis rejected the likelihood that presynaptic specializations are positioned randomly in interneurons (Fig. 5C). Our analysis also rejected the likelihood that postsynaptic specializations are positioned randomly (Fig. 5D), although the frequency distributions of distances between postsynaptic specializations are fundamentally different to those for pre-synapses (which are less numerous). These results were largely consistent for synapses in neurons grouped according to functional type or neurotransmitter type (Fig. 5E).

### Pre-synapses are characterized by varicosities but not mitochondria

In order to determine whether synapses localize to varicosities, we compared the diameter of neurons at sites of pre-synapses with the diameter at sites of post-synapses and with the average neuron diameter. In motor neurons and sensory neurons, pre-synapses, but not post-synapses, were found in the wider portions of the neurons (P=0.003 and P=0.822, respectively; Fig. 5F).

Whether synapse formation determines the site of varicosity formation, or varicosity formation determines the site of synapse formation is not known, but the co-localization of a synapse with a varicosity did not result in a difference in mitochondrial density (P=0.255; Fig. 5G).

### Mitochondria co-localize with varicosities rather than synapses

As mitochondria are most often found in varicosities (Fig. 4F), and pre-synapses also co-localize with varicosities (Fig. 5F), we tested the hypothesis that it is the synapses themselves, rather than varicosities, that are more influential in the placement of mitochondria.

Parenthetically, we did not test the possibility that mitochondria determine the distribution of varicosities (or synapses) as synapses and varicosities are stable on time scales over which mitochondria traverse long distances along these neuronal processes (Morsci et al., 2016; Mondal et al., 2021). We defined varicosities as swellings bounded on both sides by restrictions in neuronal diameter of at least 30% (Fig. 6A). We compared the proportion of synaptic varicosities containing mitochondria to the proportion of non-synaptic varicosities containing mitochondria (Fig. 6B). We found that these proportions were not statistically different in any neuron type, indicating that the presence of synapses did not predict a higher probability of an attendant mitochondrion (Fig. 6B; P=0.127 in a test of proportions across all neurons).

**Figure 6.**
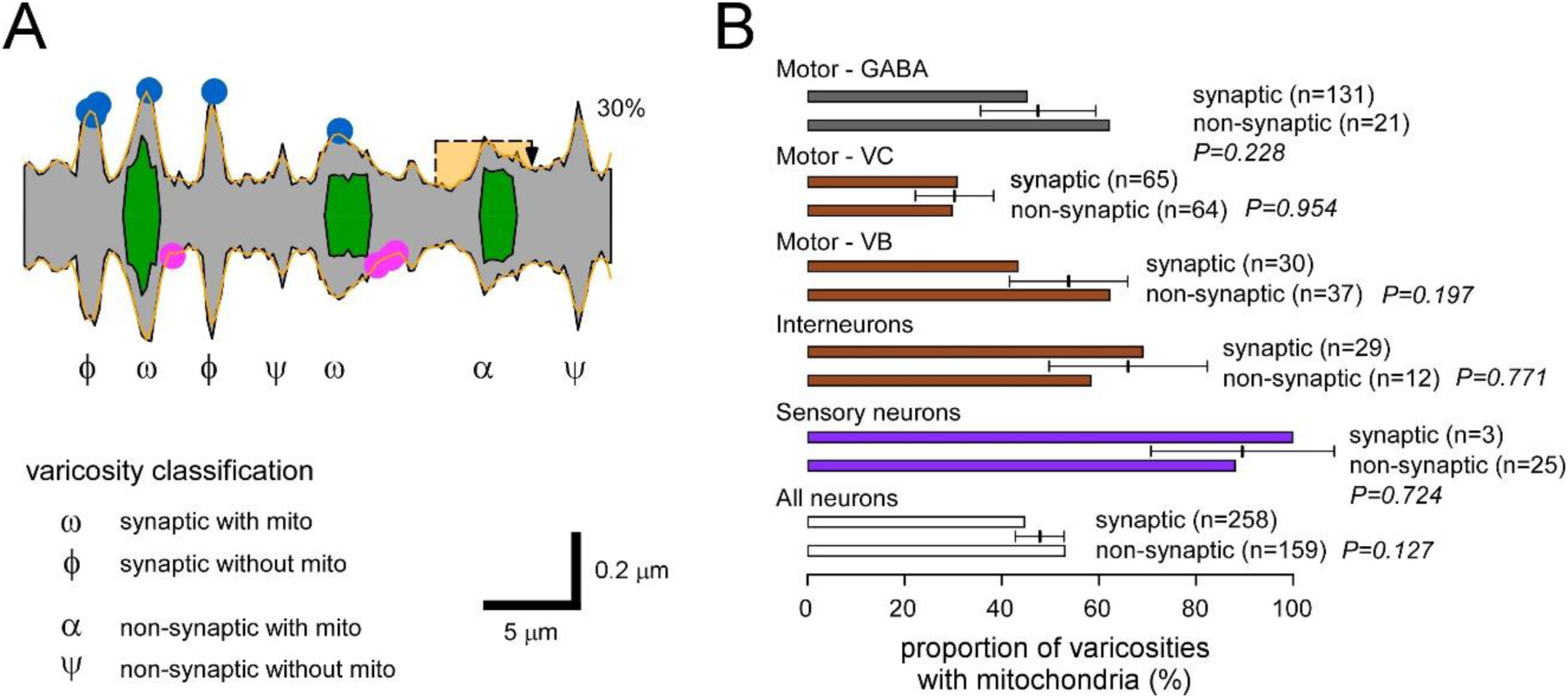
A varicosity is not more likely to contain a mitochondrion if it contains a synapse. A. The profile of a motor neuron (dorsal aspect of DD3), its mitochondria and synaptic locations, reconstructed from measurements made in each of 126 contiguous micrographs (#1588 to #1713; 30 μm) before (black trace) and after (orange trace) smoothing with a 3-point moving average. A 5 micron “hat” was placed on each swelling, and defined as a varicosity if bounded on both sides by a restricted diameter at least 30% less that the diameter of the swelling. The hat could be placed off-center, but the restrictions needed to be found with the length of the hat. Seven swellings were defined as varicosities; 4 with synapses (ω + ϕ), and 3 without (α + ψ). The proportion of synaptic varicosities bearing mitochondria can be calculated as [ω/(ω + ϕ) = 2/4 = 50%], while the proportion of non-synaptic varicosities bearing mitochondria can be calculated as [α / (α + ψ) = 1/3 = 33%]. B. A comparison of the proportion of synaptic varicosities bearing mitochondria to the proportion of non-synaptic varicosities bearing mitochondria, across neurons of different functional type. Z-tests for differences between proportions found no significant differences within any neuron type. The average pooled proportion is indicated, along with error bars (data in Suppl. Tables).

We also restricted our analyses to presynaptic varicosities, to mitigate the risk that a mitochondrial association with synaptic varicosities was being missed due to data averaging across pre- and post-synaptic compartments. Within the pre-synapse rich mixed-synapse portion of the five GABAergic motor neurons, only 45% of varicosities with a synapse (n=131) contained a mitochondrion (i.e. 55% contained no mitochondria), while 62% of varicosities without a synapse (n=21) contained a mitochondrion (P=0.228 in a test of proportions).

Similarly, an analysis restricted to pre-synapses of the three highly varicose VC-type cholinergic motor neurons yielded no sign of a mitochondrial association with synaptic varicosities (P=0.916). Previous ultrastructural studies in the hippocampus of mice and rats also reported that approximately half of all presynaptic terminals lack mitochondria (Shepherd and Harris, 1998; Chavan et al., 2015), but an important distinction made in this study is that - even when a varicosity does not contain presynaptic elements it is no less likely to contain a mitochondrion.

## DISCUSSION

For the purposes of Ca^2+^ buffering, the ready provision of ATP, or even the synthesis of neurotransmitters, a neuron might position mitochondria at synapses, yet we find little support for such control in the fine caliber neurons of *C. elegans*. In fact, our data are quite emphatic that these neurons do not systematically distribute mitochondria to synapses. To summarize, we could not dismiss the possibility that there is a random distribution of mitochondria relative to pre-synapses (Fig.4G-J). Secondly, while we could reject a random distribution of mitochondria relative to post- synapses in the most varicose neurons (Fig.4G-J), this appears to arise through the systematic exclusion of post-synapses but not mitochondria from varicosities (Fig.5F). Thirdly, mitochondria are no more likely to be found in a varicosity with a synapse, than in a varicosity without a synapse (Fig. 6). Lastly, skirting the semantics of what does or does not constitute a varicosity, mitochondrial volume density is no greater at synapses than at any other location along fine caliber neuronal processes (Fig. 5G). In light of these data, we might reexamine our assumptions about any advantage that might be gained by placing mitochondria at synapses.

In this study, the average separation between synapses and their closest mitochondrion is approximately 5μm (Fig. 4G-J), raising the question of whether this separation is likely to impair synaptic function. The Ca^2+^ microdomains of voltage-gated Ca^2+^ channels extend no more than fractions of a micron (Naraghi and Neher, 1997), and mitochondria are generally well outside this range at most synapses. Therefore, considering the rapid rate at which Ca^2+^ equilibrates over distances in the range of 1-5 μm (Sala and Hernandez-Cruz, 1990), 5 μm may still be close enough to allow mitochondria to play a role in buffering synaptic Ca^2+^, and for the Ca^2+^ to coordinate mitochondrial metabolism with synaptic activity (Chouhan et al., 2012). Similarly, a recent modelling approach predicted ATP levels to be remarkably stable in the cytosol between mitochondria, even when separated by tens of microns; albeit in the absence of inter-varicose restrictions (Kuznetsov and Kuznetsov, 2023).

Our observation of heterogeneity in mitochondrial morphometrics across neuron types (Fig. 2F-K) is consistent with previous findings in vertebrates and invertebrates (Shepherd and Harris, 1998; Cserep et al., 2018; Justs et al., 2022). Here, two findings deserve further discussion. First, we found that mitochondrial diameter is under exquisite control in the 180 to 240 nm range, to the extent that VC-type motor neurons have the capacity to deliver a diameter significantly different from the others, yet only ∼20% less (Fig. 2H). The mechanisms controlling diameter are not known, but simple restriction by the neuron itself does not appear to play a role as there is no correlation between mitochondrial diameter and neuronal diameter (Fig. S2B). We note that mitochondrial diameter correlates with neuronal diameter in the miniature parasitic wasp (Fischer et al., 2018), but neuronal diameters are as small as 98 nm in the wasp, while mitochondrial diameters are as small as 52 nm, perhaps reaching a limit of restraint if the inner membrane is to form cristae (Perkins et al., 1997; Yang et al., 2021). The second finding of interest, that mitochondrial volume density falls into one of two ranges, likely reflects different neuronal power demands. Four neuron types have a high mitochondrial volume density (5.5%, 5.9%, 6.5 and 6.5%; 6.1%±0.2% overall, mean±SEM), while one neuron type (VC-type motor neurons) is low (3.7%±0.3%, mean±SEM). These VC-type motor neurons predominantly innervate the vulval muscles, which are only intermittently active, driving muscle contraction when eggs require expulsion (Collins et al., 2016). A comparison might be made with motor neurons in *Drosophila* larvae, where type-Ib motor neurons generate consistent motor patterns driving locomotion (Lu et al., 2016; Newman et al., 2017), with an average presynaptic mitochondrial volume density of 6.3%±0.1% (mean±SEM), while another motor neuron type has a more phasic activity pattern, and a low mitochondrial density of 3.8%±0.4% (mean±SEM) (Justs et al., 2022).

On the basis of numerous ultrastructural studies we had expected to encounter sizeable differences in mitochondrial length, size and density between axons and dendrites – confirming a neuron’s capacities for subcellular control over mitochondria. However, we only observed subtle differences, limited to mitochondria with a ∼16% smaller diameter in dendrites. As *C elegans* GABAergic motor neurons do not exclude post-synapses from their axons, the neurons in our data set may provide a poor test for subcellular control. Ultrastructural studies able to substantiate a comparison of mitochondria morphometrics in axons versus dendrites of the very same neurons are quite rare because of the limited field of view in electron microscopy. Turner and others (2022) observed a clear difference in mitochondrial volume between compartments of pyramidal cells in the mouse primary visual cortex, but little to no difference was observed between axons, dendrites and somata in pyramidal cells of the mouse hippocampus (Faitg et al., 2021). Apart from emphasizing the necessity of establishing the identity of the parent cell type in electron microscopy studies comparing mitochondria across different compartments, these findings suggest that neurons may differ markedly in their capacity to control mitochondria across cellular compartments.

Finally, while it is clear that the neurons examined here do not systematically distribute mitochondria to synapses, we might also consider the possibility that invertebrates such as *C. elegans* are either unable to do so, or do not gain an advantage by doing so. Certainly, *C. elegans* has homologs of *miro*, *milton*, VAPB and FHL2 (Mercer et al., 2009; Xiong et al., 2009; Shen et al., 2016; Cottee et al., 2017), although we know of no homolog for syntaphilin. Never-the-less, there is a staggering degree of conservation of cellular mechanisms between invertebrates and vertebrates, and so it seems likely that an organism with 20,000 protein-coding genes would have the capacity for subcellular control of mitochondria if it was to its advantage. Therefore, if vertebrates truly do distribute mitochondria to synapses, while *C. elegans* does not, it would seem most plausible that *C. elegans* simply does not gain an advantage from such a design motif. Clearly, *C. elegans* are different to vertebrates by virtue of their small size alone, operating near the “small-cell limit” (Lockery and Goodman, 2009). But two features in particular might diminish a need for mitochondria at synapses; their reliance on slow neuronal action-potentials rather than regenerative all-or-none action potentials (but see (Liu et al., 2018)), and a prominent role for Ca^2+^ influx in the absence of Na^+^ channels (Goodman et al., 1998). With slower and more sustained Ca^2+^ transients in synaptic compartments, mitochondrial placement would be less critical for the purposes of either Ca^2+^ buffering or for coordinating mitochondrial energy metabolism with neuronal energy metabolism.

## MATERIALS AND METHODS

### Electron Microscopy

The electrons micrographs in *C. elegans*, examined in this manuscript, were prepared, fixed and reconstructed by White and others (White et al., 1986). Analysis was performed on digitized images of the micrographs.

### Measurement, Calibration and Analysis of Electron Micrographs

#### Cross Sectional Area Measurements

3,607 sections from the N2U section (1844-ventral portion, 1763-dorsal portion) were analyzed with the Fiji ImageJ software. Each section was 80 nm thick and centers were ∼240 nm apart (White et al., 1986; Cook et al., 2019). Cross-sectional areas of neurons and mitochondria were measured using the freehand tool. Conversion of pixel distance and area to a micron scale is described below. Cell body measurements, including those measurements on mitochondria within them, were separated from those made in the neuronal processes. Cell body measurements were those made in sections where the neuron diameter measurements of the processes exceeded 2 standard deviations of the diameter of the processes that led up to the cell body.

#### Estimation of Center-Center Distance Between Myosin Filaments

The separation between myosin filaments was calculated from images published by (Matsunaga et al., 2017) using Fiji and Matlab. Images were first processed in Fiji, utilizing the native “Thresholding” function with a range of 0-129. The results were modified with the Binary Erosion function until only the larger structures of interest remained. The “Particles” analysis was performed in order to obtain a list of all the structures and their positions in X and Y. Using Matlab, that list was then processed to calculate the distances between a point and every other point, for all points, utilizing a Euclidean distance calculation. The results were then filtered to consider only values larger than 0.04 microns and smaller than 0.06 microns. The statistics for the separation between adjacent points were thus calculated at a mean of 0.048 microns separation, with a standard deviation of 0.005 microns.

| Statistic | Value |
| --- | --- |
| Average | 0.048241 $\mu\text{m}$ |
| Standard Deviation | 0.004825 $\mu\text{m}$ |
| Median | 0.047822 $\mu\text{m}$ |
| Variance | 2.3283e-05 $\mu\text{m}^2$ |
| *N* | 345 |

#### Conversion of Pixels to Microns from Digitized Micrographs

Magnification varied between digitized electron micrographs, therefore a different factor was needed to convert pixels to microns for each micrograph. Through reference to the distance between thick filaments (48.3 nm) established above, and the number of pixels between thick filament centers on each micrograph, pixel measurements of distances and areas could be converted to μm and μm^2^, respectively, for each micrograph. A micrograph specific estimate of the pixel distance between thick filaments was made using the straight-line selection tool. We made two separate measurements of the pixel distance of 4 contiguous inter-myosin filament spacings, averaged the two, then divided by 4.

Distance and Area are therefore converted as follows:

Distance (μm) = Pixel Distance x (48.3 / inter-myosin filament pixel distance)

Area (μm^2^) = Pixel Area (Number of Pixels) x ((48.3)^2^ / (inter-myosin filament pixel distance)^2^)

Mitochondrial length was calculated by multiplying 240 nm (thickness of three sections) by the number of micrographs the mitochondria appear in, then subtracting 80 nm (see Consequences of limited axial resolution below). Mitochondrial volume was calculated by multiplying mitochondrial length by mitochondrial average area. Mitochondrial diameter was calculated by dividing mitochondrial average area by Pi, taking the square root and multiplying it by two. Instantaneous mitochondrial density was calculated by dividing mitochondrial average area by neuronal area over the same extent. Neuronal volume (includes mitochondrial volume) was calculated by multiplying average neuronal area by neuronal length. Neuronal length was calculated by multiplying 240 nm (thickness of three sections) by the number of micrographs the neuron appears in. Average mitochondrial volume density was calculated by multiplying mitochondrial average volume by the number of mitochondria present in the neuron divided by neuronal volume.

#### Consequences of limited axial resolution

The diagram below shows the range of possible lengths for a mitochondrion that appears in exactly three sections.

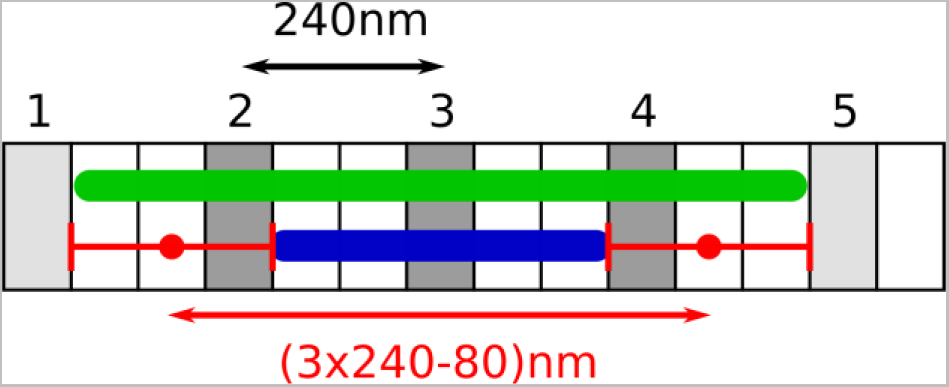

The numbered gray sections are the ones we analyzed. The green mitochondrion extends from the right edge of section 1 to the left edge of section 5, entering neither section 1 nor 5. The blue mitochondrion extends from the right edge of section 2 to the left edge of section 4, occupying just enough of sections 2 and 4 to be visible in both. The true edges of a mitochondrion could be anywhere between those two extremes. The corresponding mean locations and uncertainties are shown as red dots and red error bars, respectively. The mean mitochondrion length is 80nm shorter than the total length of the three section triplets in which the mitochondrion appears (3 x 240nm).

The uncertainty is ±240nm. Since the midpoint of the mean edge locations coincides with the midpoint of the first and last section in which the mitochondrion is seen, no correction is required when computing distances between object centers.

The diagram below illustrates another concern that should be addressed.

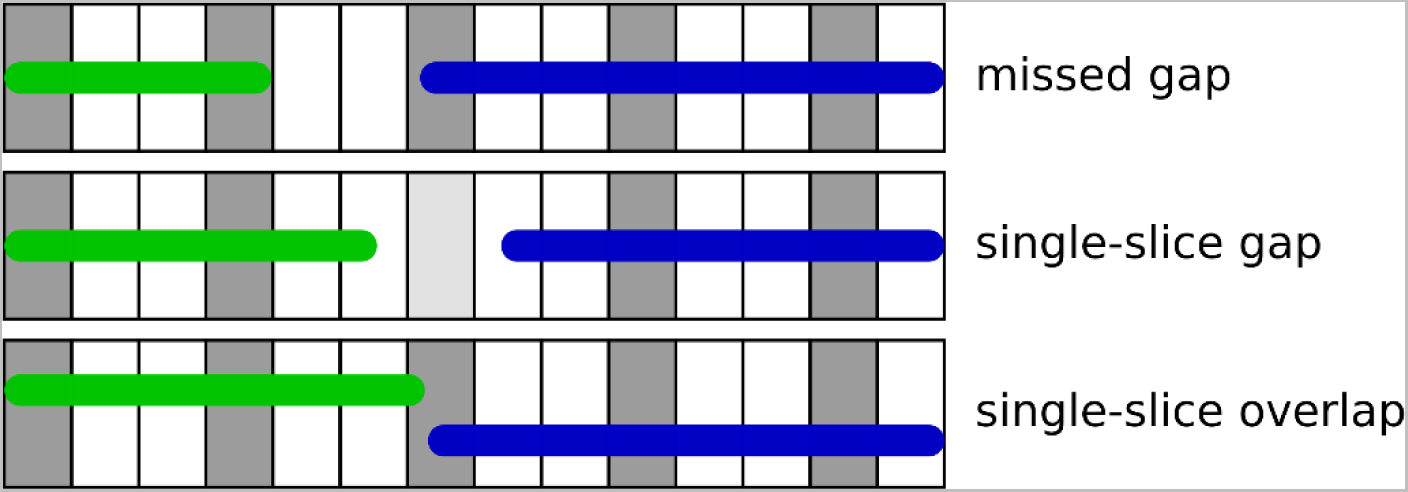

In the top scenario, there is no observable gap between the green mitochondrion and the blue mitochondrion (no empty gray section). As a result, they appear to us as a single mitochondrion. To estimate the frequency of such undetected gaps, we measure the frequency of the most similar events in the data set: single-section gaps and single-section overlaps. From a total 411 pairs of neighboring mitochondria, we find 9 single-section gap and 13 single-section overlaps. Therefore, barring a specific mechanism favoring very short gaps (0 to 80nm) while excluding slightly larger gaps (80 to 240nm) as well as short overlaps, we expect the number of missed gaps to be of the same order, about 2-3% of the sample size – not enough to significantly alter the shape of the distribution or its mean.

### Application of the Kolmogorov-Smirnov (KS) model

To test whether mitochondria are distributed randomly, we examine the distances between them. If mitochondrial centers are distributed randomly along the axon and their locations are independent of each other, then the distance between consecutive mitochondria centers should follow an exponential distribution. Similarly, if synapses are distributed randomly and independently of each other, then the distance between consecutive synapses should follow an exponential distribution. Finally, if mitochondria are distributed randomly and independently of synapses, then the distances between each synapse and the nearest mitochondrion should follow an exponential distribution.

To determine whether a distance distribution differs significantly from the exponential distribution, we used the bootstrap method and the KS statistic. First, we fit the experimental distribution of distances using the maximum likelihood estimation. The resulting exponential parameter is simply the mean distance. Second, we computed the KS statistic for the experimental distribution by computing the maximum of the difference between the experimental and fitted cumulative distribution functions. Third, we drew 10,000 random samples from the fitted distribution, with the same sample size as the experimental sample. Fourth, we computed the KS statistic for each of those 10,000 bootstrap samples. To account for the fact that exponential parameter was not known in advance, each bootstrap sample is compared with the exponential distribution whose parameter matches its own mean distance. We also accounted for the fact that distances between mitochondria or synapse centers are always multiples of half the distance between EM sections by rounding every value in the bootstrap samples to the nearest multiple of half the inter-section distance. Finally, we computed the fraction of bootstrap samples whose KS statistic was larger than the experimental sample’s KS statistic, which is the P value. This analysis was performed using the Python programming language, including the KS test provided in the scipy module (scipy.stats.kstest).

### Statistics

Student’s t-tests, ANOVA, and test for associations were performed in SigmaStat 3.5. Bonferroni’s correction was used to adjust the level of α when multiple t-tests were applied. ANOVA was performed when multiple comparisons were applied and an overall α of <0.05 was required for significance. Interactions could not be tested where data points were missing, e.g. no instances of overlapping mitochondria in VB-type MNs (Fig. 4F), and insufficient data for presynaptic and postsynaptic mitochondrial density in sensory neurons (Fig.5G). Pearson product-moment correlation coefficient was calculated to test the strength of associations. Tests for differences in proportion were performed using a z-test in SigmaStat 3.5, with Yates correction.

### Synapse Annotation

#### Motor Neurons

DD1 – GABA: Ventral cord motor neurons, reciprocal inhibitors, change synaptic pattern during L1 DD2 – GABA: Ventral cord motor neurons, reciprocal inhibitors, change synaptic pattern during L1 DD3 – GABA: Ventral cord motor neurons, reciprocal inhibitors, change synaptic pattern during L1 VD5 – GABA: Ventral cord motor neuron, innervates vent body muscles, reciprocal inhibitor.

VD4 – GABA: Ventral cord motor neuron, innervates vent body muscles, reciprocal inhibitor. VB4 – Cholinergic: Ventral cord motor neuron, innervates ventral body muscles.

VB5 – Cholinergic: Ventral cord motor neuron, innervates ventral body muscles.

VC1 – Cholinergic: Hermaphrodite specific ventral cord motor neuron innervates vulval muscles and ventral body muscles.

VC2 – Cholinergic: Hermaphrodite specific ventral cord motor neuron innervates vulval muscles and ventral body muscles.

VC3 – Cholinergic: Hermaphrodite specific ventral cord motor neuron innervates vulval muscles and ventral body muscles.

#### Interneurons – Cholinergic

AVAL- Ventral cord command interneuron

AVBL - Ventral cord command interneuron

AVAR- Ventral cord command interneuron

AVBR- Ventral cord command interneuron

#### Sensory Neurons – Glutamate

AVM- Anterior ventral microtubule cell, touch receptor

PVM- Posterior ventral microtubule cell, touch receptor

More information can be found on Wormatlas.org.

### Blender Renderings

Blender was used to render neurons in silhouette in Figures 1B and 3C. Using Blender’s Python Application Programming Interface (API), calibrated radius, depth, and position data were imported into Blender (http://www.blender.org; RRID: SCR_008606). In rare instances, where electron micrographs were missing, calibrated radius values were “patched” by assigning an average of the two immediately adjacent values in place of the missing value. As a part of the python code, cylinders were populated with the radius and depth corresponding to each section. The depth of each cylinder represents the distance between micrographs (240 nm) but these were adjusted in some instances to observe longer lengths of neuron (refer to scale bars in each figure). To represent any synapses present in each section a set of different colored objects were created and assigned accordingly, where blue represents a presynaptic terminal, pink represents a postsynaptic terminal, and yellow represents both a presynaptic and postsynaptic terminal. The python code used to import the data, create the cylinders, and assign the corresponding materials can be found at: https://github.com/scrill2015/scrill2015/blob/de3fbc6afc4ec83d74fbdb8f48e322d71afc46a9/Modeling%20C.%20elegans%20Neurons%20in%20Blender%20with%20Python%20API

## Acknowledgements

This work was supported by NIH NINDS awards NS103906 and NS123377 to GTM and NIH R24 OD010943 to DHH. We are grateful for discussions with Drs Guy Benian and Ivanov Stoyan. We thank John White and Jonathan Hodgkin for sending the collected TEM images of the MRC/LMB lab of Sydney Brenner to the Hall lab for long term conservation and sharing on WormImage (also supported by NIH R24 OD010943).

---

**Supplemental Figure 1:**
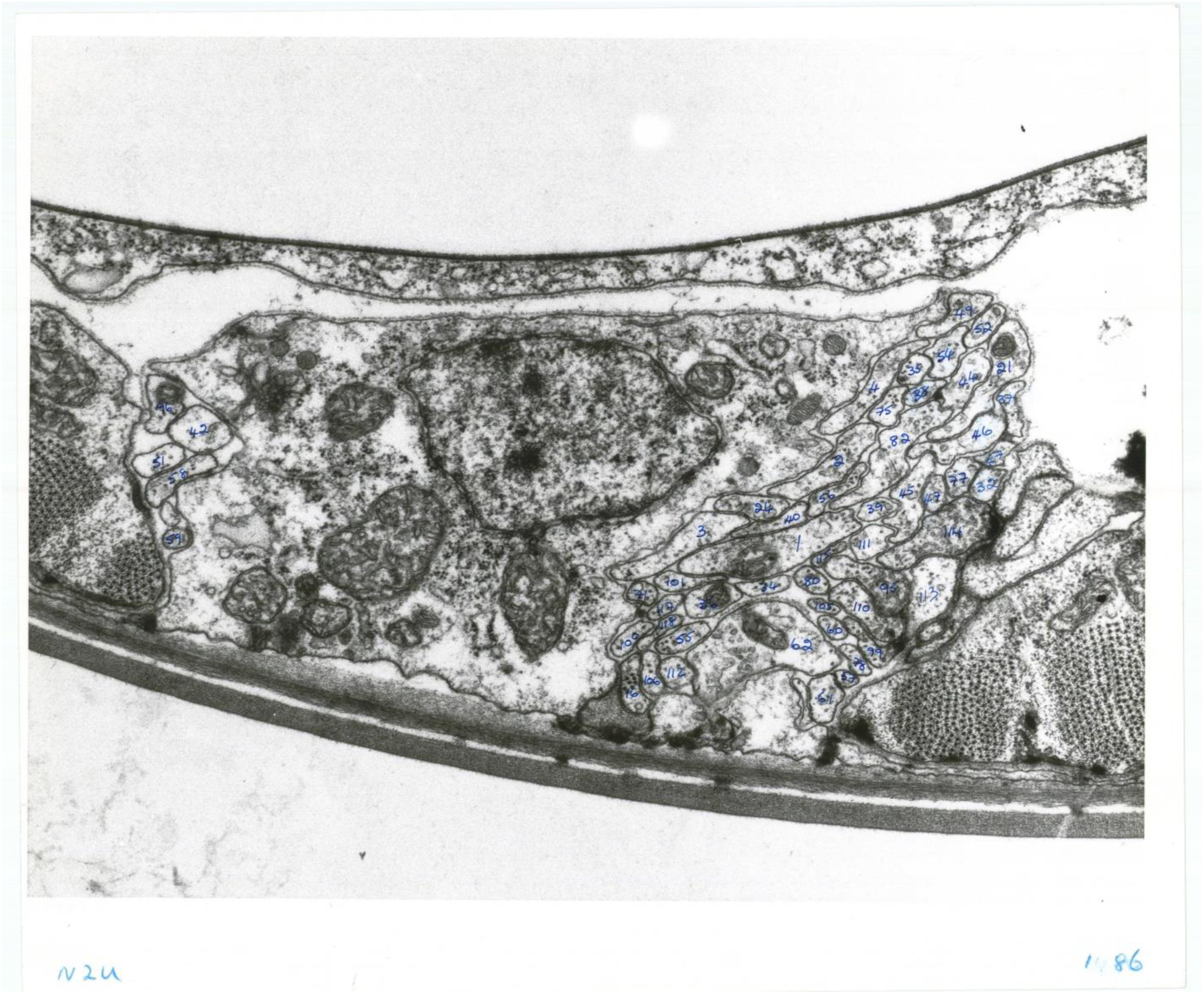
An example micrograph (N2U series #1486) from the ventral side of the worm with the central mass on the worm (intestine/gonads) missing.

**Supplemental Figure 2:**
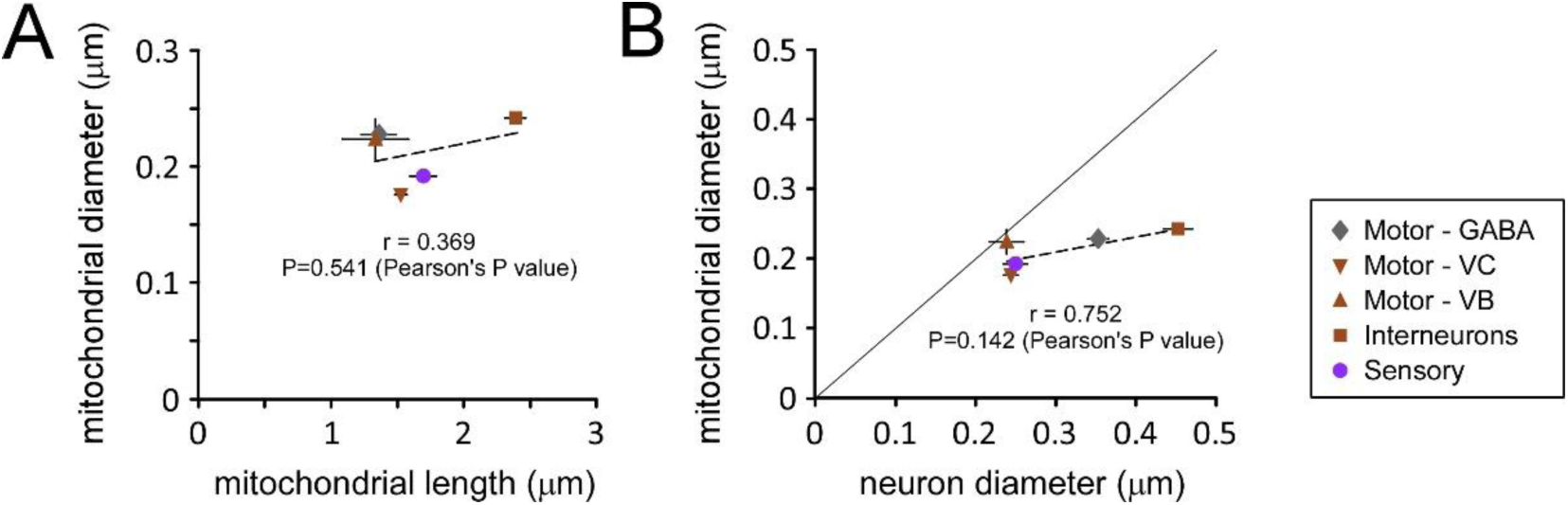
A. The regression of mitochondrial diameter on neuron diameter showed no correlation (r=0.369, P=0.541), which would be expected if neuronal diameter is not a key determinant of mitochondrial diameter. B. The regression of mitochondrial diameter on mitochondrial length also showed no correlation (r=0.752, P=0.142). Pearson’s correlation coefficients were calculated in both cases.

**Supplemental Figure 3:**
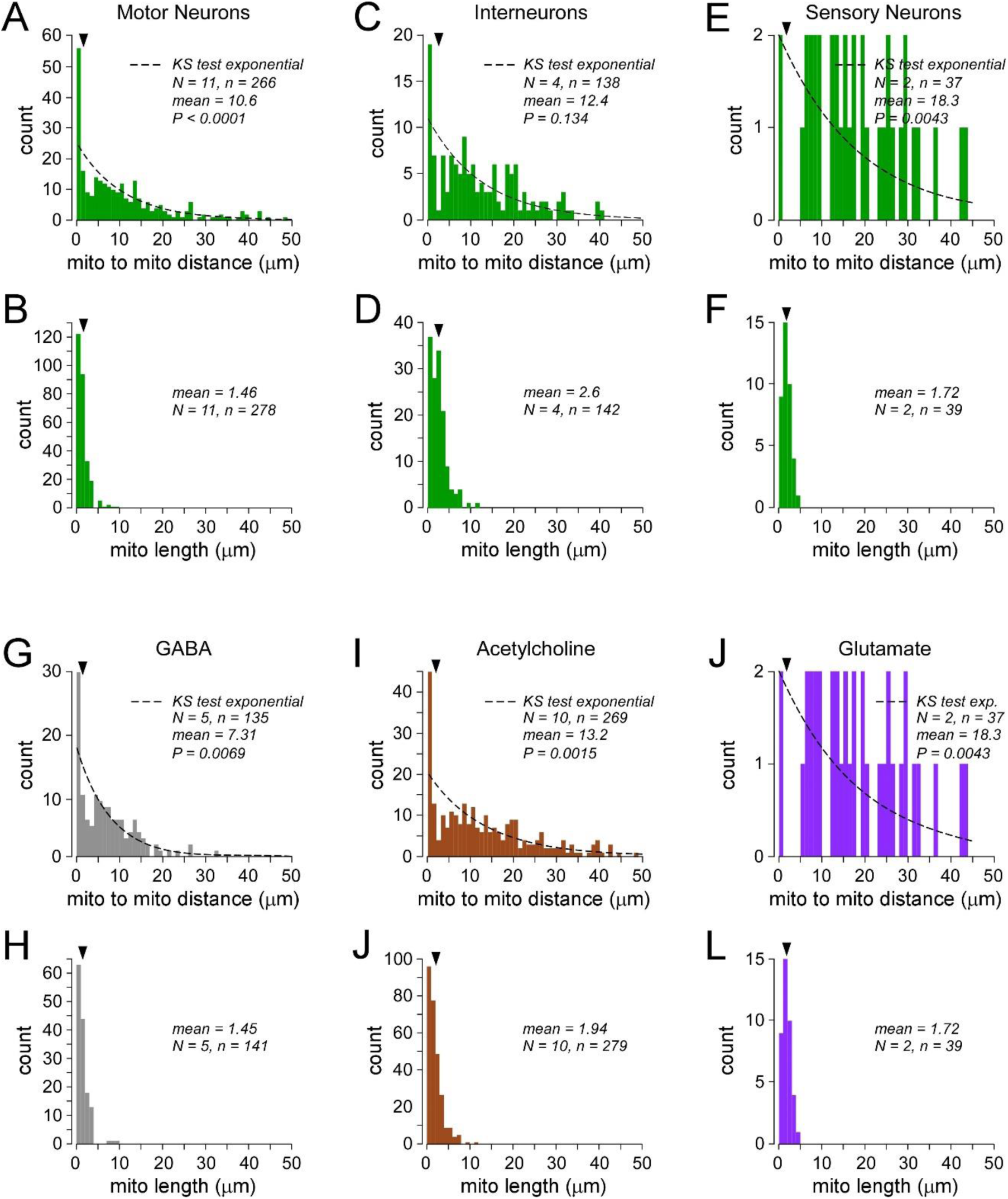
Frequency distributions of distances between mitochondria for each grouping of neurons (A, C, E, G, I and J), and the frequency distributions of mitochondrial lengths in each neuron group (B, D, F, H, J and L). Mitochondria in cell bodies were included. Results of KS tests against random distributions are shown for each interval plot. The mean mitochondrial length was calculated for each neuron grouping and shown (arrowhead) above the frequency distributions of both mitochondrial lengths and the distances between those mitochondria.

## REFERENCES

Bapat O, Purimetla T, Kruessel S, Shah M, Fan R, Thum C, Rupprecht F, Langer JD, Rangaraju V (2024) VAP spatially stabilizes dendritic mitochondria to locally support synaptic plasticity. Nat Commun 15:205.

Basu H, Pekkurnaz G, Falk J, Wei W, Chin M, Steen J, Schwarz TL (2021) FHL2 anchors mitochondria to actin and adapts mitochondrial dynamics to glucose supply. J Cell Biol 220.

Chavan V, Willis J, Walker SK, Clark HR, Liu X, Fox MA, Srivastava S, Mukherjee K (2015) Central presynaptic terminals are enriched in ATP but the majority lack mitochondria. PLoS One 10:e0125185.

Cheng XT, Huang N, Sheng ZH (2022) Programming axonal mitochondrial maintenance and bioenergetics in neurodegeneration and regeneration. Neuron 110:1899–1923.

Chouhan AK, Ivannikov MV, Lu Z, Sugimori M, Llinas RR, Macleod GT (2012) Cytosolic calcium coordinates mitochondrial energy metabolism with presynaptic activity. J Neurosci 32:1233–1243.

Collins KM, Bode A, Fernandez RW, Tanis JE, Brewer JC, Creamer MS, Koelle MR (2016) Activity of the C. elegans egg-laying behavior circuit is controlled by competing activation and feedback inhibition. Elife 5.

Cook SJ, Jarrell TA, Brittin CA, Wang Y, Bloniarz AE, Yakovlev MA, Nguyen KCQ, Tang LT, Bayer EA, Duerr JS, Bulow HE, Hobert O, Hall DH, Emmons SW (2019) Whole-animal connectomes of both Caenorhabditis elegans sexes. Nature 571:63–71.

Cottee PA, Cole T, Schultz J, Hoang HD, Vibbert J, Han SM, Miller MA (2017) The C. elegans VAPB homolog VPR-1 is a permissive signal for gonad development. Development 144:2187–2199.

Cserep C, Posfai B, Schwarcz AD, Denes A (2018) Mitochondrial Ultrastructure Is Coupled to Synaptic Performance at Axonal Release Sites. eNeuro 5.

Devine MJ, Kittler JT (2018) Mitochondria at the neuronal presynapse in health and disease. Nat Rev Neurosci 19:63–80.

Faitg J, Lacefield C, Davey T, White K, Laws R, Kosmidis S, Reeve AK, Kandel ER, Vincent AE, Picard M (2021) 3D neuronal mitochondrial morphology in axons, dendrites, and somata of the aging mouse hippocampus. Cell Rep 36:109509.

Fischer S, Lu Z, Meinertzhagen IA (2018) From two to three dimensions: The importance of the third dimension for evaluating the limits to neuronal miniaturization in insects. J Comp Neurol 526:653–662.

Goodman MB, Hall DH, Avery L, Lockery SR (1998) Active currents regulate sensitivity and dynamic range in C. elegans neurons. Neuron 20:763–772.

Hammond JW, Huang CF, Kaech S, Jacobson C, Banker G, Verhey KJ (2010) Posttranslational modifications of tubulin and the polarized transport of kinesin-1 in neurons. Mol Biol Cell 21:572–583.

Huang B, Jones SA, Brandenburg B, Zhuang X (2008) Whole-cell 3D STORM reveals interactions between cellular structures with nanometer-scale resolution. Nat Methods 5:1047–1052.

Justs KA, Lu Z, Chouhan AK, Borycz JA, Lu Z, Meinertzhagen IA, Macleod GT (2022) Presynaptic Mitochondrial Volume and Packing Density Scale with Presynaptic Power Demand. J Neurosci 42:954–967.

Kang JS, Tian JH, Pan PY, Zald P, Li C, Deng C, Sheng ZH (2008) Docking of axonal mitochondria by syntaphilin controls their mobility and affects short-term facilitation. Cell 132:137–148.

Kruppa AJ, Buss F (2021) Motor proteins at the mitochondria-cytoskeleton interface. J Cell Sci 134.

Kuznetsov IA, Kuznetsov AV (2023) ATP diffusional gradients are sufficient to maintain bioenergetic homeostasis in synaptic boutons lacking mitochondria. Int J Numer Method Biomed Eng 39:e3696.

Lebiedzinska M, Szabadkai G, Jones AW, Duszynski J, Wieckowski MR (2009) Interactions between the endoplasmic reticulum, mitochondria, plasma membrane and other subcellular organelles. Int J Biochem Cell Biol 41:1805–1816.

Liu Q, Kidd PB, Dobosiewicz M, Bargmann CI (2018) C. elegans AWA Olfactory Neurons Fire Calcium-Mediated All-or-None Action Potentials. Cell 175:57–70 e17.

Lockery SR, Goodman MB (2009) The quest for action potentials in C. elegans neurons hits a plateau. Nat Neurosci 12:377–378.

Lopez-Domenech G, Kittler JT (2023) Mitochondrial regulation of local supply of energy in neurons. Curr Opin Neurobiol 81:102747.

Lu Z, Chouhan AK, Borycz JA, Lu Z, Rossano AJ, Brain KL, Zhou Y, Meinertzhagen IA, Macleod GT (2016) High-Probability Neurotransmitter Release Sites Represent an Energy-Efficient Design. Curr Biol 26:2562–2571.

Matsunaga Y, Hwang H, Franke B, Williams R, Penley M, Qadota H, Yi H, Morran LT, Lu H, Mayans O, Benian GM (2017) Twitchin kinase inhibits muscle activity. Mol Biol Cell 28:1591–1600.

Mercer KB, Szlam SM, Manning E, Gernert KM, Walthall WW, Benian GM, Gutekunst CA (2009) A C. elegans homolog of huntingtin-associated protein 1 is expressed in chemosensory neurons and in a number of other somatic cell types. J Mol Neurosci 37:37–49.

Mondal S, Dubey J, Awasthi A, Sure GR, Vasudevan A, Koushika SP (2021) Tracking Mitochondrial Density and Positioning along a Growing Neuronal Process in Individual C. elegans Neuron Using a Long-Term Growth and Imaging Microfluidic Device. eNeuro 8.

Morsci NS, Hall DH, Driscoll M, Sheng ZH (2016) Age-Related Phasic Patterns of Mitochondrial Maintenance in Adult Caenorhabditis elegans Neurons. J Neurosci 36:1373–1385.

Naraghi M, Neher E (1997) Linearized buffered Ca2+ diffusion in microdomains and its implications for calculation of [Ca2+] at the mouth of a calcium channel. J Neurosci 17:6961–6973.

Newman ZL, Hoagland A, Aghi K, Worden K, Levy SL, Son JH, Lee LP, Isacoff EY (2017) Input-Specific Plasticity and Homeostasis at the Drosophila Larval Neuromuscular Junction. Neuron 93:1388–1404 e1310.

Perkins G, Renken C, Martone ME, Young SJ, Ellisman M, Frey T (1997) Electron tomography of neuronal mitochondria: three-dimensional structure and organization of cristae and membrane contacts. J Struct Biol 119:260–272.

Sala F, Hernandez-Cruz A (1990) Calcium diffusion modeling in a spherical neuron. Relevance of buffering properties. Biophys J 57:313–324.

Shen Y, Ng LF, Low NP, Hagen T, Gruber J, Inoue T (2016) C. elegans miro-1 Mutation Reduces the Amount of Mitochondria and Extends Life Span. PLoS One 11:e0153233.

Shepherd GM, Harris KM (1998) Three-dimensional structure and composition of CA3-->CA1 axons in rat hippocampal slices: implications for presynaptic connectivity and compartmentalization. J Neurosci 18:8300–8310.

Sigal YM, Speer CM, Babcock HP, Zhuang X (2015) Mapping Synaptic Input Fields of Neurons with Super-Resolution Imaging. Cell 163:493–505.

Turner NL et al. (2022) Reconstruction of neocortex: Organelles, compartments, cells, circuits, and activity. Cell 185:1082–1100 e1024.

Wang X, Schwarz TL (2009) The mechanism of Ca2+ -dependent regulation of kinesin-mediated mitochondrial motility. Cell 136:163–174.

White JG, Southgate E, Thomson JN, Brenner S (1986) The Structure of the Nervous-System of the Nematode Caenorhabditis-Elegans. Philosophical Transactions of the Royal Society B-Biological Sciences 314:1–340.

Winding M, Pedigo BD, Barnes CL, Patsolic HG, Park Y, Kazimiers T, Fushiki A, Andrade IV, Khandelwal A, Valdes-Aleman J, Li F, Randel N, Barsotti E, Correia A, Fetter RD, Hartenstein V, Priebe CE, Vogelstein JT, Cardona A, Zlatic M (2023) The connectome of an insect brain. Science 379:eadd9330.

Xiong G, Qadota H, Mercer KB, McGaha LA, Oberhauser AF, Benian GM (2009) A LIM-9 (FHL)/SCPL-1 (SCP) complex interacts with the C-terminal protein kinase regions of UNC-89 (obscurin) in Caenorhabditis elegans muscle. J Mol Biol 386:976–988.

Yang Z, Wang L, Yang C, Pu S, Guo Z, Wu Q, Zhou Z, Zhao H (2021) Mitochondrial Membrane Remodeling. Front Bioeng Biotechnol 9:786806.

